# Anatomy-Guided 3D Graph Networks for Couinaud Segmentation in Tumor Affected Livers

**DOI:** 10.64898/2026.05.11.724316

**Authors:** Lei You, Haotian Dang, Hongyu Wang, Eduardo J Matta, Xiaobo Zhou

## Abstract

Image-based liver Couinaud segmentation is designed to automatically provide the locations of suspicious objects in liver CT/MR images. Once achieved, the physicians will be guided to the target slice and area where the suspicious node is located. However, conventional algorithms trained primarily on healthy liver images often fail to generalize to Hepatocellular Carcinoma (**HCC**) cases due to pathological structural distortions. In this work, we propose a robust two-stage framework that integrates a 3D Unet with a 3D Anatomical Structure-Guided Graph Convolutional Network (3D GCN). This two-stage strategy effectively isolates the liver volume to eliminate structural noise from neighboring organs, such as the spleen, allowing the framework to focus exclusively on the complex 3D anatomical relationships among the eight segments. To ensure the topological consistency required for global spatial reasoning, we implement a standardized preprocessing pipeline that normalizes liver-only volumes to exactly 50 frames along the z-axis. By combining a lightweight 3D UNet backbone with the 3D GCN for refined boundary reasoning, our model demonstrates superior generalization performance on unseen clinical datasets, achieving a mean Dice score of 0.828 in blind testing. By releasing our code and pretrained weights, we aim to provide the first publicly available deep learning resource for robust Couinaud segmentation.

## Introduction

Precise delineation of hepatic anatomy is fundamental to the diagnosis, treatment planning, and surgical management of liver diseases, particularly hepatocellular carcinoma (HCC), in which tumor burden, vascular invasion, and parenchymal remodeling frequently distort normal structural landmarks. The Couinaud classification, the most widely adopted framework for functional liver anatomy, divides the liver into eight functionally independent segments, each characterized by its own vascular inflow, biliary drainage, and venous outflow [1-3]. Within each segment, a central branch of the portal vein, hepatic artery, and bile duct supplies the parenchyma, whereas the hepatic veins course between segments, forming the natural intersegmental boundaries used across radiology, surgery, and interventional oncology [2, 3]. This functional organization provides a universal language for clinical communication and supports accurate segment-level localization, which is essential for lesion characterization, surgical resection, ablative therapy planning, and longitudinal disease monitoring [3, 4].

Despite its centrality in clinical workflows, reliable Couinaud segmentation from routine CT and MRI remains challenging. Traditional image-based algorithms perform adequately in healthy livers but often fail to generalize pathological conditions, where distorted vasculature, necrotic tumor regions, and altered parenchyma obscure anatomical boundaries. Many models are trained predominantly on healthy anatomy, resulting in limited robustness when faced with the heterogeneity typical of HCC or cirrhosis. This generalization gap limits their applicability in real-world oncologic settings, underscoring the need for segmentation frameworks that preserve anatomical fidelity across diverse clinical presentations.

Recent advances in deep learning have substantially improved organ and suborgan segmentation accuracy; however, the most robust performance gains have come from approaches that explicitly incorporate anatomical priors and structural dependencies into the learning process. To address the limitations of prior methods, we introduce an anatomy-guided, two-stage computational framework that integrates a 3D U-Net with a 3D anatomical structure-guided Graph Convolutional Network (3D-GCN). This framework leverages a high-performance teacher network trained on a heterogeneous dataset comprising both healthy and HCC-affected livers, improving generalizability across a wide range of anatomical variations. To support efficient clinical deployment, we apply knowledge distillation using a composite loss function that combines Kullback–Leibler divergence and Dice normalization, enabling the distilled student model to maintain high segmentation accuracy at reduced computational cost.

The framework was trained using publicly available datasets (MSD08 and LiTS) and rigorously validated on internal clinical data, demonstrating consistently strong performance across all eight Couinaud segments. The distilled student model achieves an average Dice score of 0.889, closely matching the teacher model’s Dice of 0.890, highlighting the effectiveness of the distillation strategy. By releasing our code and pretrained weights, we provide the first publicly available deep learning resource capable of robust Couinaud segmentation in both healthy and HCC-affected livers, offering a scalable solution that advances liver cancer imaging research and enhances clinical decision support.

## Related work

Deep learning models have demonstrated significant achievements in medical tasks, such as organ segmentation[5], disease classification[6], data generation[7] and so on[8, 9]. Deep learning approaches for Couinaud liver segmentation have evolved along several distinct branches. Most prevalent are fully convolutional segmentation methods, which leverage U-Net-like architectures and skip connections to classify each voxel into one of nine anatomical segments. These models can be implemented as either 2.5D slice-based[10] or fully 3D frameworks[11, 12]. However, given the prohibitive computational demands of high-resolution 3D processing, researchers frequently down-sample input volumes to lower resolutions (e.g., 256 × 256 × 64). To improve feature representations, Liang Qiu et al[13] stacked multiple transformer layers at the bottleneck of the U-Net structure. The multi-head attention block of the transformer layers helps to refine the global features by calculating the QKV relationships among feature vectors. Dong Miao et al[14] attempt to address the Couinaud segmentation task by incorporating landmarks to improve the segmentation results of their 3D UNet.

There are many false positives near the boundaries, such as segments 6/7 and segments 5/8, due to the lack of clear physical landmarks and the high degree of inter-individual variability in the course of the right hepatic vein and the branching patterns of the right portal vein. Zhang et al try to solve this by learning complementary feature representations from Couinaud points and voxels. Qian et al[15] combine the cloud points from liver, vessel and boundary information to enhance the feature representation. Tian et al[10] proposed an Attention-Guided Reasoning (ARR-GCN) framework to explicitly model the anatomical spatial relationships between liver segments. In their graph formulation, nodes represent the feature embeddings of the eight anatomical segments, while edges encode their prior adjacency as defined by a learnable matrix. Through graph convolution, these node features are updated to capture global structural context and subsequently projected back to refine the backbone UNet’s pixel-wise predictions. However, the study does not extensively evaluate robustness in pathological scenarios, where tumors often distort these anatomical boundaries. Furthermore, the reliance on a 2.5D input approach (e.g., 7-slice stacks) inherently restricts volumetric reasoning, which may compromise segmentation performance when generalizing to unseen 3D CT data with variable slice thickness.

In this paper, we propose a robust two-stage framework that integrates a lightweight 3D U-Net with a novel Dynamic 3D Anatomical Structure-Guided Graph Convolutional Network (3D-AS-GCN). Our innovations include: 1. Our hierarchical two-stage strategy first localizes the liver to constrain the search space, allowing the subsequent GCN to focus entirely on resolving internal segmental boundaries. Based on the first-stage segmentation, we build a 3D data-preprocessing pipeline to handle the variable number of slices along the Z-axis. 2. We develop the dynamic 3D graph reasoning mechanism, which moves beyond standard pixel-wise classification by explicitly modeling anatomical connectivity in 3D space. This module uses a soft-weighted average to aggregate voxel-level features into discrete anatomical nodes and employs a depth-adaptive adjacency matrix that interpolates connectivity rules along the z-axis. 3. To facilitate reproducibility and further research, we will publicly release all source code and pretrained model weights.

## Methods

As shown in **Figure1**, there are three stages for obtaining the Couinaud liver segments from the input 3D .nii files. A 3D nnUNet [16] is fine-tuned to obtain coarse liver and tumor segmentation masks in the first stage. Using those masks, we extract the liver regions, referred to as liver-only scans, from abdominal CT scans and normalize them to a resolution of 512 × 512×50. In the final stage, we use the liver-only scans to train the 3D GCN, which performs the Couinaud segmentation. Evaluation matrices include the Dice score and Average Symmetric Surface Distance (ASSD) for each segment, as well as the mean values across all segments.

**Figure 1.**
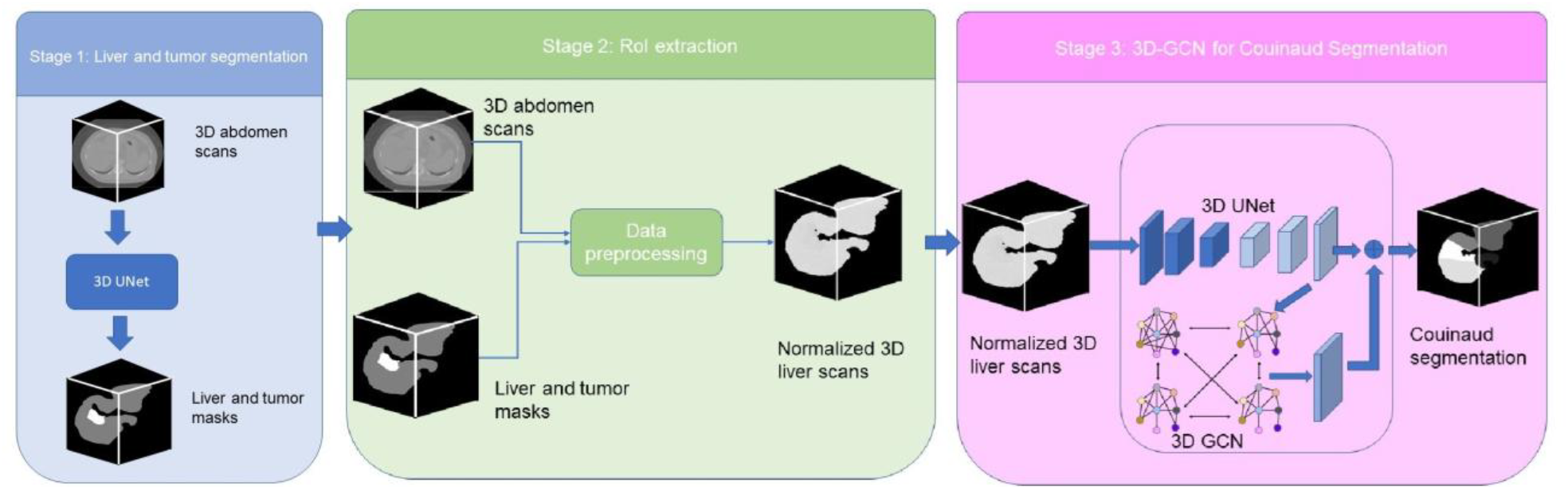
The architecture of our liver Couinaud segmentation network.

### Liver and tumor segmentation

In our two-stage model, we employed the standard, self-adapting nnUNet model for liver segmentation. We trained two separate nnUNet models to perform 1) liver-tumor mask segmentation, 2) Couinaud mask segmentation as the baseline model. We adhere to the default nnUNet-v2 configuration, the network self-adapting topology, preprocessing steps, and five-fold split training pipeline.

The nnUNet pipeline first generates a CT data fingerprint based on trainset’s statistics F = Φ(*D*), where *D* denotes the data itself. The fingerprint *F* contains voxel spacing, image shapes, and CT intensity percentiles. Based on the data fingerprint and computational hardware configuration *H*, the nnUNet planner generates the training plan *P* :

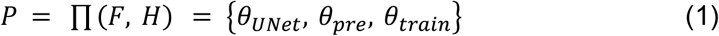

where *θ*_*UNet*_ represents the self-adapted topology of nnUNet, *θ*_*pre*_ represent the preprocessing rules of CT image, *θ*_*train*_ represents training parameters. During training, the loss function adopted by nnUNet is always:

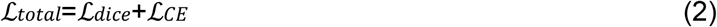

where ℒ_*CE*_ represent voxel cross-entropy loss, and dice loss is defined by,

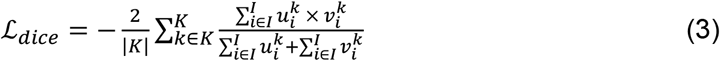

where *u* represents the softmax probability output of the network, *v* represents a one hot encoding of ground truth segmentation map, *i* represents the number of pixels, and *k* represents the classes.

The models were trained from scratch and went through all folds’ training cross-validation schemes to ensure robustness. During inference, the nnUNet utilizes the plan configuration to preprocess input data and infer the masks.

### Couinaud Segmentation with 3D GCN

To represent the 3D Couinaud liver anatomy as a graph, our model utilizes a soft-aggregation mechanism to convert volumetric voxel features into discrete anatomical nodes. For an input feature map 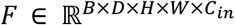 from our 3DUNet, where *Cin* represents the channel dimension of the latent features, we aggregate spatial information using a corresponding coarse probability map *P* ∈ [0,1]^*B*×*D*×*H*×*W*×*K*^. This probability map is derived from the softmax output of the same U-Net, where K represents the 8 liver segments plus background. For each batch b, slice depth d, and segment class *i* ∈ {1, …., *K*}, the node feature vector *n*_*b,d,i*_ ∈ *R*^*Cin*^ is defined as the weighted average of voxel features based on their class probabilities.

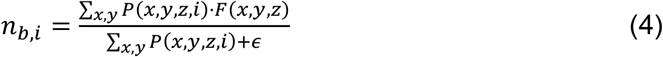

where *P*(*x, y, z, i*) represents the possibility that a voxel at spatial coordinates (x,y,z) belongs to segment i, and *F*(*x, y, z*) is the corresponding features at that location. We use a smaller depth d instead of the original z to optimize GPU memory allocation during training and inference.

We use *H* = {*n*_*b,i*_ | *b* ∈ [1, *B*], *i* ∈ [1, *K*]|} ∈ ℝ^*K*×*Cin*^ as the eight 3D node features and a precalculated adjacent matrix A as the edges of the 3D GCN. In the original GCN literature[17], the network is defined as following.

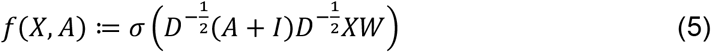

In our case, we modify the equation to get an updated feature matrix *H*^*l*+1^ from *H*^*l*^ with the following equation.

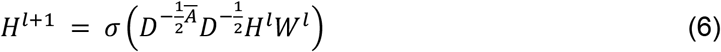

Where *Ā* = *A* + *I* and D is used to perform a symmetric normalization of *Ā*. W is the learned weights of the 3DGCN. Each diagonal element of the degree matrix D represents the sum of the edge weights for a specific node as shown in the following equation.

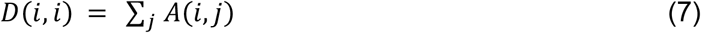

The refined anatomical information is projected back into the voxel space by computing the dot product between the enhanced node features *H*^*l*+1^ and the original voxel-level feature map F. This operation calculates a class-specific influence for every voxel based on the global anatomical context captured by the nodes. It creates a residual correction map, P_refined_, where each voxel’s value represents an anatomically reasoned “vote” for a specific segment.

The P_refined_ is directly added to the coarse output P_coarse_ produced by the 3D U-Net backbone, expressed as P_final_ = P_coarse_ + P_refined_. This operation incorporates prior anatomical knowledge into the original network’s output prior to the final activation step. The primary benefit of this hybrid architecture is that the final segmentation mask becomes significantly more robust to the structural distortions and atypical intensities characteristic of pathological scans.

## Experiment and results

### Datasets

The LiTS2017 dataset comprises 131 CT scans with a mean voxel size of 0.793 × 0.793 × 1.506 mm. The average slice thickness for this cohort is 1.506 mm (SD: 1.173 mm). For this dataset, Couinaud segment annotations were obtained from established literature[18]. The MSD08 dataset provided 193 available Couinaud annotations derived from literature[19]. The MSD08 volumes exhibit a mean voxel size of 0.821×0.82×4.563 mm, with a significantly larger mean slice thickness of 4.563 mm (SD: 1.419 mm). Our private dataset comprises 500 CT scans and has a mean voxel size of 0.898×0.898×2.477 mm. The average slice thickness for this cohort is 2.477 mm (SD: 0.871 mm). We manually annotated 100 cases for the generalization evaluation of our 3D GCN. Additionally, we used the PLC-CECT liver CT dataset from J.Luo et al[20] in our liver segmentation model training. It consists of 278 patients with 1112 CT images. The voxel size for this cohort is uniformly 1×1×1 mm.

All data files were preprocessed into liver-specific volumes with an initial resolution of 512×512×50 by our liver segmentation network. To accommodate GPU memory constraints and maintain the integrity of the 3D GCN anatomical prior, all stage-two volumes were resampled to a standardized depth of 32 slices, and restricted to the regions containing the liver. This normalization ensures that the spatial relationships within the 3D adjacency matrix remain topologically consistent across a diverse clinical population. With a resolution of 8×8×32, the adjacency matrix effectively models the global connectivity of the liver volume.

### Model implementation details

In our experiment, we use the Pytorch 2.8.0 with CUDA 12 as the default environment for our networks. The models are trained with a Nvidia V100 GPU with 32 GB of memory. To accelerate the convergence of the 3D GCN, the 3D UNet backbone was pre-trained for 75 epochs; its weights were then used to initialize the 3D GCN, which was trained for an additional 100 epochs. The models are trained with a batch size of 4 and a learning rate of 1e-4 using the Adam optimizer. The nnUNet model training adopted the nnUNetv2 3d_fullres default setup and planner It is trained for 1000 epochs, with 250 minibatches, an SGD optimizer, and a learning rate that descends from 0.01 to 1e-5.

The architecture of our 3D UNet backbone follows a classical encoder-decoder structure with four levels of down-sampling and up-sampling blocks. The encoder progressively doubles the feature depth from 64 to 1024 channels at the bottleneck, while the decoder utilizes skip connections to preserve high-resolution spatial information. Notably, the model is configured to produce a dual-resolution output: a low-resolution feature map (f_lowres) and a full-resolution feature map (f_fullres), enabling hierarchical integration of local and global anatomical details.

The 3D GCN architecture comprises two sequential graph convolutional layers that refine the feature embeddings extracted by the 3D UNet backbone. The module accepts input features with a dimension of 128 (C_in=128). The first GCN layer reduces this to a hidden dimension of 64 with a ReLU activation, while the second layer restores the dimension to 128 to facilitate residual learning. By aggregating voxel-level embeddings into node-level representations, the network utilizes the 3D adjacency matrix to reason over anatomical structures. These refined nodes are then redistributed back to the voxel space, yielding a segmentation mask that strictly adheres to known anatomical priors.

Evaluation and matrix: We use a five-fold validation method to assess segmentation performance by computing the mean Dice score and ASSD. The Dice score is a spatial overlap metric that measures the similarity between the predicted segmentation (P) and the ground truth (G).

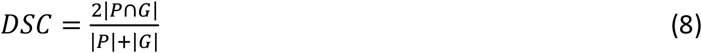

While Dice focuses on volume, **ASSD** focuses on the alignment of the boundaries (*S*_*P*_ and *S*_*G*_). It calculates the average distance between all points on the surface of the prediction and the nearest points on the surface of the ground truth, and vice versa.

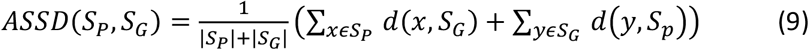

Where *S*_*P*_and *S*_*G*_ are the sets of surface points for the prediction and ground truth respectively. The function d calculates the shortest Euclidean distance from point x/y to the surface *S*_*G*_/*S*_*P*_.

To demonstrate the robustness and generalization of our proposed framework, we conducted a comparative analysis against several state-of-the-art methods in liver Couinaud segmentation. As shown in Table 1, while several existing studies rely exclusively on a single data source—such as the private-only cohorts used by Tingting Xie[11] (0.94 Dice) and Liang Qiu[13] (0.82 Dice)— our model maintains consistently high performance across three diverse datasets, achieving 0.8362 on LiTS2017, 0.8444 on MSD08, and 0.828 on our private validation set. Furthermore, our approach demonstrates superior cross-dataset generalization compared to multi-benchmark evaluations, such as those by Jin Qian[15], who reported 0.8164 and 0.8253 on LiTS and MSD, respectively. This stability is primarily attributed to our two-stage refinement strategy, which yielded a significant 9.5% improvement in mean Dice score over the single-stage baseline as shown in Table 2. Unlike many competing models that utilize fixed data splits, our results are grounded in a rigorous five-fold cross-validation protocol, ensuring that our performance metrics are representative and highly reliable for clinical applications.

**Table 1.** recent studies related to CT-based Couinaud segmentation

| teams | MSD08 | LiTS2017 | Private data | Sample size | Data split | Mean Dice |
| --- | --- | --- | --- | --- | --- | --- |
| Liang Qiu [13] | no | no | yes | 123 | 87:18:18 | 0.82 |
| Xukun Zhang [12] | Yes | Yes | no | 191(MSD)<br>131(LiTS) | 95:29:67<br>63:20:48 | 0.8946(MSD)<br>0.9145(LiTS) |
| Tingting Xie [11] | no | no | yes | 348 | 170:178 | 0.94 |
| Fei Lyu [21] | yes | no | no | 161 | 113:16:32 | 0.7352 |
| Yinli Tian [10] | yes | no | no | 112 | 92:20 | 0.895 |
| Seungyoo Lee [22] | no | no | yes | 193 | 115:28:50 | 0.8417 |
| Jin Qian [15] | yes | yes | no | 191(MSD)<br>131(LiTS) | 7:2:1 | 0.8253(MSD)<br>0.8164(LiTS) |
| **Ours** | yes | yes | yes | 191(MSD)<br>131(LiTS)<br>100(Private, validation only) | Five-fold | 0.8444(MSD)<br>0.8362(LiTS)<br>0.828(Private data) |

**Table 2.** The mean dice score of our two-stage model (LiTS2017+MSD08).

|  | Fold | Mean Dice | Seg 1 | Seg 2 | Seg 3 | Seg 4 | Seg 5 | Seg 6 | Seg 7 | Seg 8 |
| --- | --- | --- | --- | --- | --- | --- | --- | --- | --- | --- |
| single stage | 1 | 0.7501 | 0.7608 | 0.7442 | 0.7160 | 0.7983 | 0.7293 | 0.7311 | 0.7618 | 0.7592 |
|  | 2 | 0.7722 | 0.7791 | 0.8206 | 0.7570 | 0.7975 | 0.7226 | 0.7390 | 0.8276 | 0.7346 |
|  | 3 | 0.8079 | 0.8199 | 0.8129 | 0.8166 | 0.8326 | 0.7434 | 0.7796 | 0.8598 | 0.7983 |
|  | 4 | 0.765 | 0.8098 | 0.7601 | 0.7738 | 0.8202 | 0.7692 | 0.7591 | 0.7211 | 0.7067 |
|  | 5 | 0.6697 | 0.7859 | 0.4063 | 0.2344 | 0.7794 | 0.7509 | 0.7406 | 0.8410 | 0.8190 |
|  | Average | 0.753 | 0.7911 | 0.7088 | 0.6595 | 0.8056 | 0.7430 | 0.7498 | 0.8023 | 0.7636 |
| two stages | 1 | 0.8489 | 0.8661 | 0.8732 | 0.8549 | 0.8732 | 0.7961 | 0.796 | 0.8839 | 0.8481 |
|  | 2 | 0.8588 | 0.872 | 0.9019 | 0.8617 | 0.8554 | 0.8164 | 0.8332 | 0.8831 | 0.8467 |
|  | 3 | 0.8615 | 0.869 | 0.8894 | 0.8647 | 0.8676 | 0.8314 | 0.8251 | 0.8889 | 0.8563 |
|  | 4 | 0.839 | 0.8518 | 0.8698 | 0.8526 | 0.8521 | 0.7995 | 0.7935 | 0.8629 | 0.8302 |
|  | 5 | 0.8311 | 0.8547 | 0.861 | 0.7915 | 0.8486 | 0.8018 | 0.8023 | 0.8616 | 0.8276 |
|  | Average | **0.8479** | **0.8627** | **0.879** | **0.845** | **0.8594** | **0.809** | **0.81** | **0.8761** | **0.8418** |

### Ablation study

#### Two-stage segmentation

In this section, we evaluate whether a two-stage segmentation strategy improves Couinaud liver segmentation compared to a single-stage approach, while maintaining consistency in the training datasets and network architecture. For the data used in the single-stage model, we kept the original z-axis dimensions while resampling the xy resolution to 256×256. From these volumes, we extracted patches of 256×256×32 for model training. These patches further enabled the dynamic computation of a 3D adjacency matrix that captures the spatial co-occurrence and topological relationships among liver segments across the training cohort.

As shown in Table 2, the two-stage strategy consistently achieves a higher Dice score across all eight segments than the single-stage approach, with a mean Dice of 0.847. For adjacent regions such as Segment 5/8 and Segment 6/7, the two-stage strategy yields average increases of approximately 7.2% and 6.7%, respectively, demonstrating that isolating the liver volume effectively eliminates structural interference from irrelevant neighboring organs such as the spleen. By removing this anatomical noise, the 3D GCN can focus its feature aggregation on the complex 3D relationships and adjacency constraints among the eight internal liver segments, rather than processing non-target tissues, resulting in more precise boundary refinement for early liver cancer screening.

#### GCN reasoning

As shown in Table 3, the integration of 3D GCN reasoning provides a substantial performance gain, with the Mean Dice score increasing from 0.7311 for the backbone model to 0.8241. The 3D GCN model outperforms the backbone in nearly all individual segments, effectively capturing complex 3D anatomical information. While Segment 1 has the most anatomical connections with other segments, the backbone model fails to distinguish it from surrounding tissues, yielding a low Dice score of 0.1609. In contrast, the 3D GCN provides a robust and stable segmentation result with a score of 0.8143. Both models perform well on the adjacent segments of the right lobe. Segments 6 and 7 are predicted with high accuracy, with the 3D GCN further refining these scores to 0.8236 and 0.8639, respectively. For adjacent segments 5 and 8, both models maintain predictive power above 0.76, though GCN reasoning specifically improves Segment 8’s accuracy to 0.8215.

**Table 3.** the improvements brought by 3D GCN reasoning on our private data

|  | Mean Dice | Seg 1 | Seg 2 | Seg 3 | Seg 4 | Seg 5 | Seg 6 | Seg 7 | Seg 8 |
| --- | --- | --- | --- | --- | --- | --- | --- | --- | --- |
| 3D UNet(backbone) | 0.7311 | 0.1609 | 0.8496 | 0.8372 | 0.7608 | 0.7708 | 0.8069 | 0.8495 | 0.8138 |
| 3D GCN | 0.8241 | 0.8143 | 0.8646 | 0.8596 | 0.7767 | 0.7684 | 0.8236 | 0.8639 | 0.8215 |

#### Compared with a baseline model

we evaluate Couinaud segmentation performance by comparing our 3D GCN model against the nnUNet baseline across several public datasets. As shown in Table 4, nnUNet achieves higher scores in the five-fold cross-validation for each dataset. However, it is important to note the disparity in computational requirements: the nnUNet was trained using default settings for 1,000 epochs with a dynamic learning rate. In contrast, our 3D GCN requires significantly less training, consisting of 75 backbone pre-training epochs and 100 GCN refinement epochs. Furthermore, our model maintains a much smaller memory footprint (300 MB) compared to nnUNet (500 MB). Despite this lightweight architecture and reduced training time, our 3D GCN delivers competitive performance.

**Table 4.** Liver Couinaud segmentation results on public datasets with fivefold validation

|  | 3D GCN |  |  |  | nnUNet |  |  |  |
| --- | --- | --- | --- | --- | --- | --- | --- | --- |
|  | MSD08 |  | LiTS2017 |  | MSD08 |  | LiTS2017 |  |
| Fold | Mean Dice | Mean ASSD | Mean Dice | Mean ASSD | Mean Dice | Mean ASSD | Mean Dice | Mean ASSD |
| 1 | 0.8461 | 0.5903 | 0.8418 | 0.6064 | 0.8821 | 1.6813 | 0.8681 | 6.8263 |
| 2 | 0.8489 | 2.6268 | 0.8283 | 0.7374 | 0.8884 | 1.5272 | 0.8727 | 5.5333 |
| 3 | 0.8559 | 1.4119 | 0.8442 | 0.5326 | 0.8831 | 1.6083 | 0.8626 | 6.6552 |
| 4 | 0.8318 | 1.6569 | 0.8128 | 0.6945 | 0.8985 | 1.7340 | 0.8177 | 7.5993 |
| 5 | 0.8396 | 1.6487 | 0.8536 | 0.4635 | 0.8739 | 1.7578 | 0.8467 | 5.4489 |
| Average | 0.8444 | **1.5869** | 0.8362 | **0.6069** | **0.8852** | 1.6617 | **0.8535** | 6.4126 |

Most notably, our 3D GCN demonstrates superior boundary accuracy as measured by the ASSD score. On the LiTS2017 dataset, our model achieves a mean ASSD of 0.6069 mm, a significant improvement over the 6.4126 mm recorded by nnUNet. A similar advantage is observed on the MSD08 dataset, where the 3D GCN yields a mean ASSD of 1.5869 mm, outperforming the baseline score of 1.6617 mm. These results indicate that the integration of 3D anatomical priors enables more precise surface delineation and boundary adherence across diverse clinical volumes, even with a more efficient architectural design.

Notably, our 3D GCN model demonstrates superior generalization capabilities on external datasets. In this experiment, both models were trained on a composite dataset of LiTS2017 and MSD08, then evaluated on a cohort of 100 cases from our clinical center. As shown in Table 5, our model outperforms nnUNet by 12% on this private clinical dataset. While we observed a marginal decrease in mean Dice scores compared to the LiTS2017 and MSD08 benchmarks, this is attributed to the increased complexity of our local data, which encompasses diverse scan protocols and multi-vendor hardware variability.

**Table 5.**
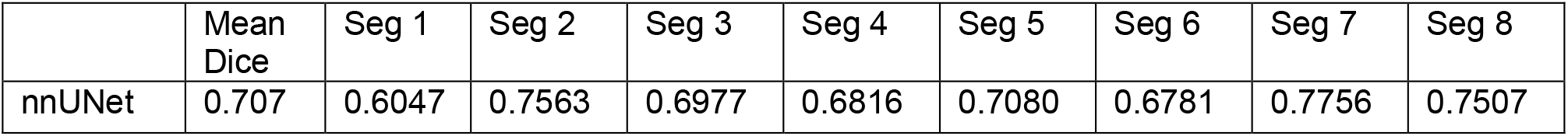
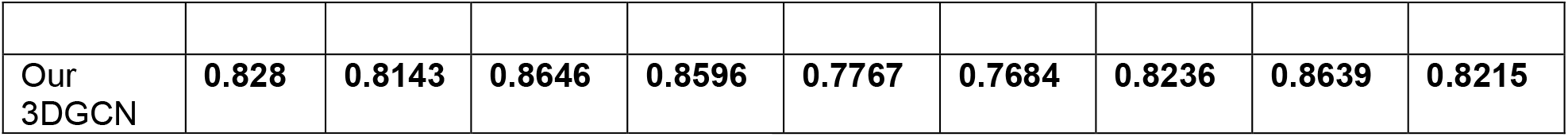
The Liver Couinaud Segmentation Results on Private Dataset with Pretrained Models

In **Figure 2**, we present a comparative visualization of the segmentation results for nnUNet and our 3D GCN against the ground truth (GT) across six private clinical cases. Representative axial slices were selected to highlight the challenges of delineating adjacent segments where anatomical boundaries are easily confused. Notably, in slice z=20 of Case 2, nnUNet fails to accurately segment segments 5, 6, 7, and 8, whereas our 3D GCN produces refined results that closely align with the GT. A similar trend is observed in slice z=25 of Case 3 and Case 4, where the 3D GCN maintains structural integrity despite the complex adjacency among the segments. These qualitative visualizations directly support the quantitative findings in Table 5, where our 3D GCN demonstrates superior generalization performance across the private dataset, particularly in segments 5 through 8, significantly outperforming the nnUNet baseline in mean Dice score.

**Figure 2.**
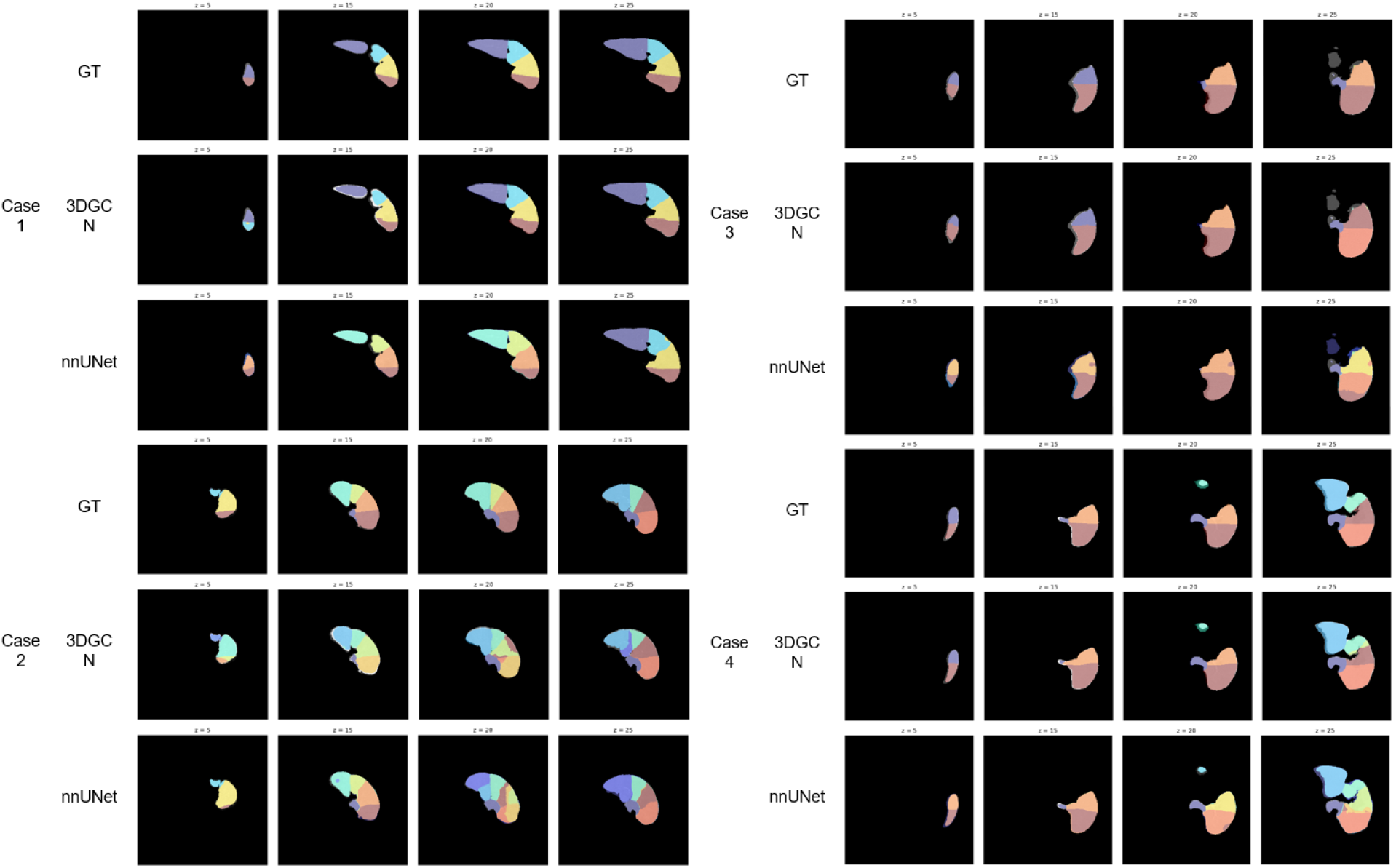
We show the differences among the GT annotations, 3DGCN segmentations, and nnUNet segmentations across different slices along z axis.

## Discussion

While deep learning has advanced Couinaud liver segmentation, achieving clinical-grade reliability remains a significant hurdle due to the extreme heterogeneity of CT data across different medical institutions. A primary challenge stems from the contrasting pathological presentations in public benchmarks: the LiTS2017 dataset features a higher density of smaller lesions (averaging 6.93 per case, each 11.25 mL), whereas the MSD08 dataset is characterized by fewer but significantly larger tumors (averaging 1.80 per case, each 77.68 mL). Since Couinaud segments are defined by the underlying topology of the hepatic and portal veins, validating models on tumor-heavy datasets such as MSD08 rigorously tests their ability to infer anatomical boundaries under severe vessel distortion. To address this, we evaluated a 3D GCN model trained on a fusion of LiTS2017 and MSD08 on a private clinical dataset comprising cirrhosis and HCC patients, achieving a Mean Dice score of 0.828. This generalization performance is further contextualized by the UMAP latent feature analysis in **Figure 3**, which reveals distinct, non-overlapping clusters for the LiTS2017 (yellow) and MSD08 (green) datasets. Notably, the private data points (black) are widely dispersed across the entire feature space, covering and extending beyond both public clusters. This distribution confirms that while real-world clinical practice encompasses a much broader spectrum of liver conditions and scan protocols than public datasets alone, our 3D GCN effectively utilizes anatomical priors to maintain robust performance across unseen clinical domains.

**Figure 3.**
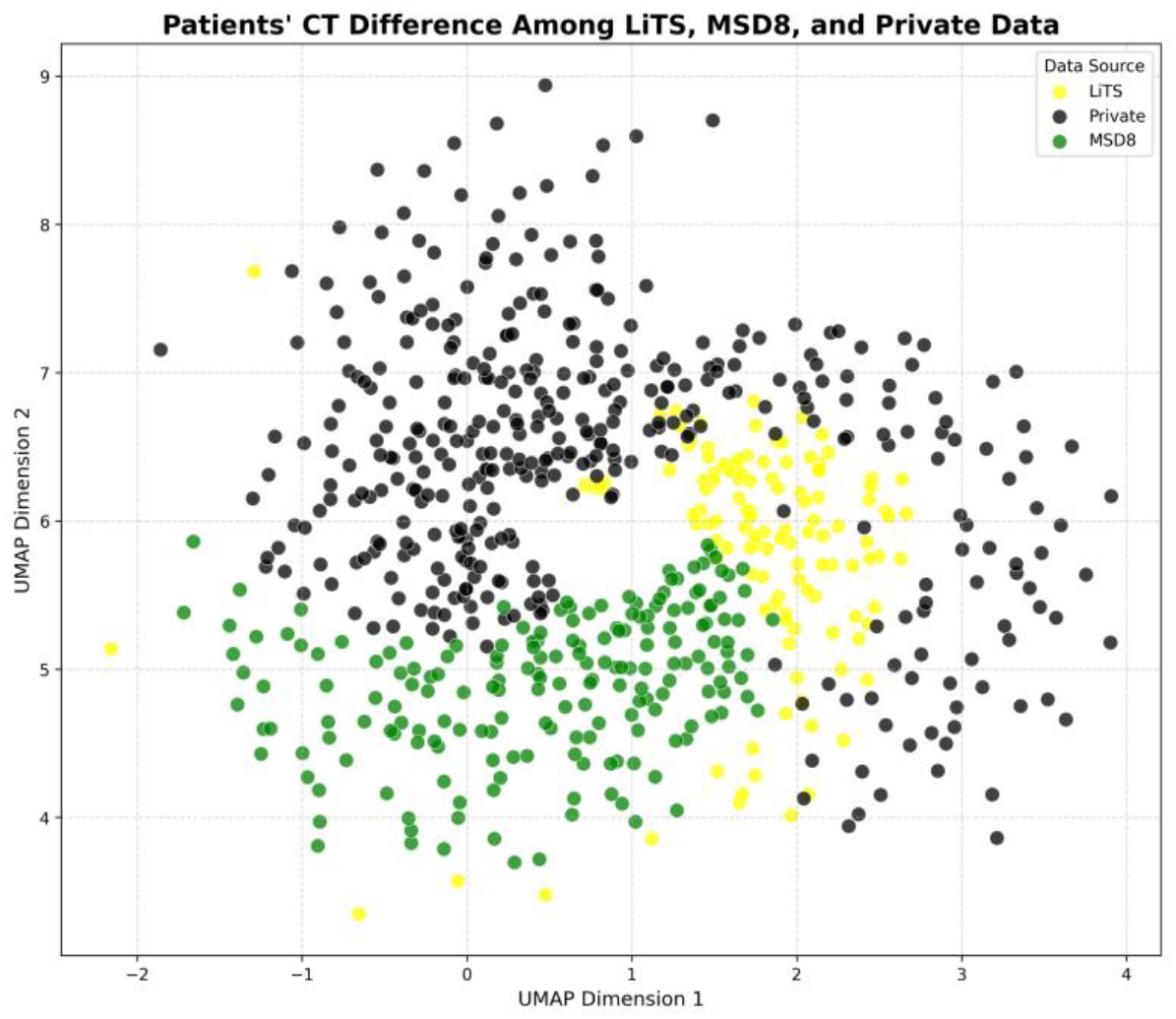
The distribution of three datasets, where yellow points, green points, and black points represent the LiTS2017, MSD08, and our private data, respectively.

A significant challenge in current research is the limited availability of open-source code and pretrained model weights, which restricts the objective validation of new frameworks across independent cohorts. Furthermore, inconsistent data partitioning across recent studies—such as the 95:29:67 split used by Xukun Zhang (approximately a 10:3:7 ratio)[12] or the 92:20 ratio used by Yinli Tian[10] —often results in models that exhibit poor generalization on unseen third-party datasets. To address these discrepancies, we evaluated our 3D GCN framework through comprehensive ablation studies anchored by a robust five-fold cross-validation protocol. Our results highlight the stability of the proposed model across diverse experimental conditions, specifically demonstrating high performance in blind tests on our private clinical dataset, achieving a mean Dice score of 0.828. To facilitate future advancements and transparency in Couinaud liver segmentation, we will release our full source code and pretrained weights to the research community upon the official publication of this work.

## Conclusion

In this study, we proposed a lightweight network for Couinaud segmentation that integrates a 3D GCN with a 3D UNet backbone. By leveraging a 3D adjacency matrix to refine latent features, our model achieves state-of-the-art results on public datasets. Furthermore, the architecture demonstrates robust generalization on unseen clinical data, maintaining performance across diverse scan protocols and multi-vendor hardware variability. We intend to release the source code and model weights upon publication to support further research.

In conclusion, while our two-stage 3D GCN framework achieved a robust mean Dice score of 0.828 on our private data, significantly outperforming the nnUNet baseline (0.707), we identified specific technical scenarios that warrant further refinement. As illustrated in **Figure 4**, several “Hard” cases with low Dice scores, such as 0.295, were found in close proximity to “Easy” cases with high scores, such as 0.970, suggesting that the model occasionally fails to differentiate between failure cases and those with similar data representations. These failures are often rooted in initial segmentation errors, where false positives from adjacent organs such as the spleen or discontinuities caused by large, invasive tumors degrade subsequent Couinaud refinement. Furthermore, the current global 3D positional reasoning is challenged by post-operative anatomical variations, such as in patients who have undergone major hepatic resections and no longer possess the standard eight-segment structure. To address these constraints, our future work will focus on developing a joint liver-and-tumor segmentation framework and an adaptive, localized 3D reasoning model capable of accommodating varied clinical presentations and post-operative anatomy.

**Figure 4.**
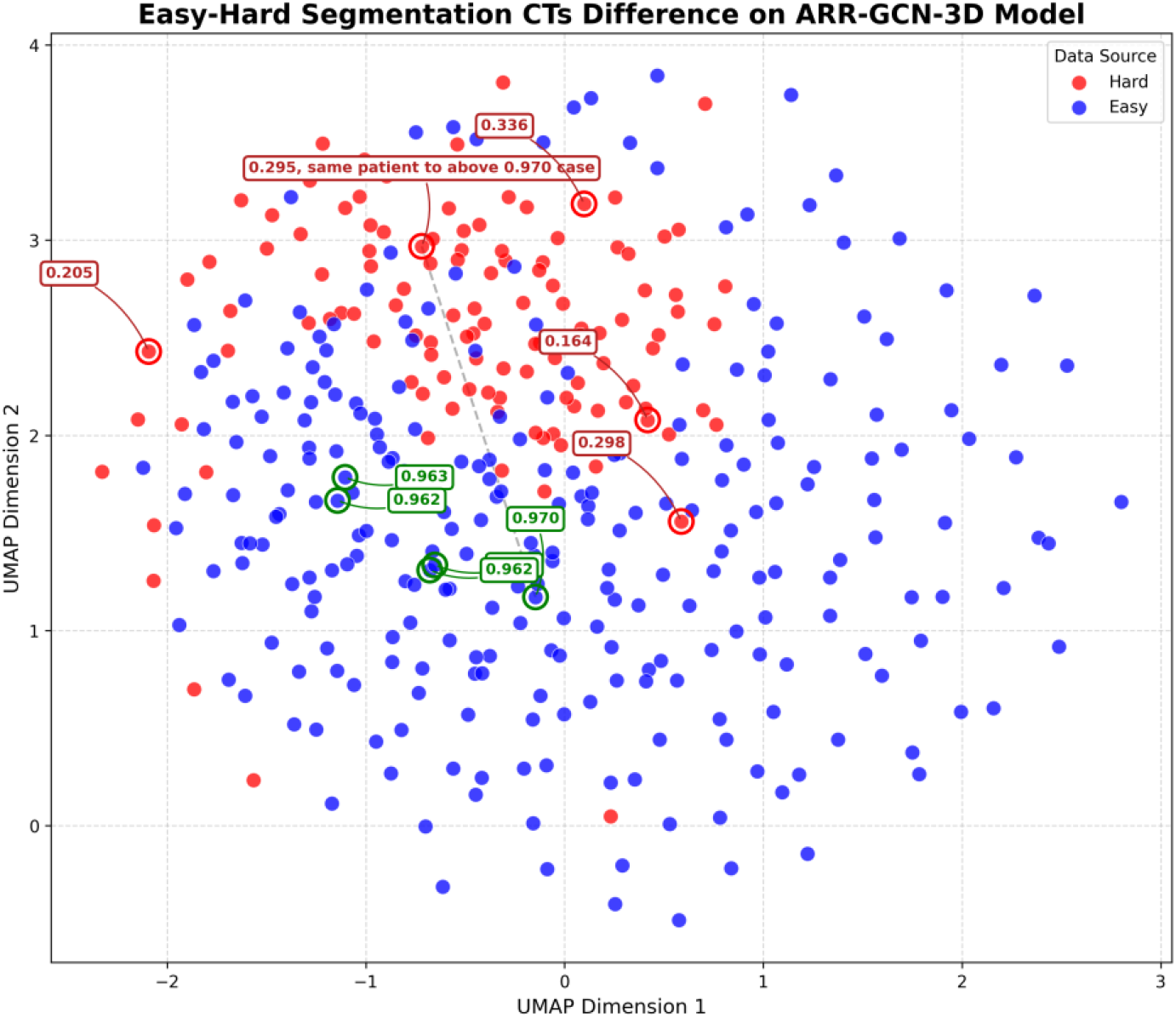
We show the difference between easy and hard CT segmentation cases (with a threshold of 0.7 for segmentation DSC similarity to Ground Truth) in our private dataset.

## Code availability

We will release our source codes at https://github.com/lei-you/Anatomy-Guided-3D-Graph-Network-for-Couinaud-Segmentation-in-Tumor-Affected-Livers.

## Acknowledgement

This project is supported by the CPRIT-funded project (GRANT ID RP250043).

